# Shallow whole genome sequencing for robust copy number profiling of formalin-fixed paraffin-embedded breast cancers

**DOI:** 10.1101/231480

**Authors:** Suet Feung Chin, Angela Santoja, Marta Grzelak, Soomin Ahn, Stephen-John Sammut, Harry Clifford, Oscar M. Rueda, Michelle Pugh, Mae A. Goldgraben, Helen A. Bardwell, Eun Yoon Cho, Elena Provenzano, Federico Rojo, Emilio Alba, Carlos Caldas

**Affiliations:** Cancer Research UK Cambridge Institute, Li Ka Shing Centre, University of Cambridge, Robinson Way, Cambridge CB2 0RE, UK; Department of Oncology, University of Cambridge, Cambridge CB2 2QQ, UK; Medical Oncology Service, Hospital Universitario Regional y Virgen de la Victoria, Instituto de Investigación Biomédica de Málaga (IBIMA), Málaga, Spain & Laboratorio de Biología Molecular del Cáncer, Centro de Investigaciones Médico-Sanitarias (CIMES), Universidad de Málaga, Málaga, Spain; Department of Pathology, Seoul National University Bundang Hospital, 82, Gumi-ro 173 Beon-gil, Bundang-gu, Seongnam, Gyeonggi, 13620, Korea. Inivata, Li Ka Shing Centre, Robinson Way, Cambridge CB2 0RE, UK; Inivata UK, The Portway Building, Granta Park, Cambridge, CB21 6GS, United Kingdom; Department of Medical Genetics, University of Cambridge, Cambridge CB2 0QQ; Department of Pathology and Translational Genomics, Samsung Medical Center, Sungkyunkwan University School of Medicine, 50 Irwon-dong, Gangnam-gu, Seoul, 135-710, South Korea; Cambridge Breast Unit, Addenbrooke’s Hospital, Cambridge University Hospital NHS Foundation Trust and NIHR Cambridge Biomedical Research Centre, Cambridge CB2 2QQ, UK; Cancer Research UK Cambridge Cancer Centre, Cambridge, CB2 0QQ, UK; Pathology Department, Instituto de Investigación Sanitaria Fundación Jiménez Díaz (IIS-FJD), Madrid, Spain.; GEICAM-Spanish Breast Cancer Research Group, Madrid, Spain

**Author notes:** Co-corresponding author &.

**Keywords:** Formalin-fixed paraffin-embedded (FFPE), shallow whole genome sequencing (sWGS), copy number (CN) and breast cancer

## Abstract

Pathology archives with linked clinical data are an invaluable resource for translational research, with the limitation that most cancer samples are formalin-fixed paraffin-embedded (FFPE) tissues. Therefore, FFPE tissues are an important resource for genomic profiling studies but are under-utilised due to the low amount and quality of extracted nucleic acids. We profiled the copy number landscape of 356 breast cancer patients using DNA extracted FFPE tissues by shallow whole genome sequencing. We generated a total of 491 sequencing libraries from 2 kits and obtained data from 98.4% of libraries with 86.4% being of good quality. We generated libraries from as low as 3.8ng of input DNA and found that the success was independent of input DNA amount and quality, processing site and age of the fixed tissues. Since copy number alterations (CNA) play a major role in breast cancer, it is imperative that we are able to use FFPE archives and we have shown in this study that sWGS is a robust method to do such profiling.

---

Comparative Genomic Hybridisation (CGH) ^1^ has had a significant impact in the study of cancer genomes. Chromosomal regions gained or lost in the tumour could be easily visualised by hybridization onto normal human metaphase spreads, allowing characterisation of genome-wide copy number alterations (CNA) in tumours ^1^. Microarrays with DNA probes (cloned DNA or oligonucleotides) spotted onto glass slides representing the entire genome soon replaced normal chromosomes ^2^ making it faster and easier to profile. The importance of characterizing somatic CNAs in cancer is now well established, with a recent TCGA pan-cancer analysis showing that human tumours can be classified into mutation driven (M-class) or copy-number driven (C-class) subtypes. Breast cancer is a C-class cancer type ^3^ and we have previously shown that CNAs are the main determinants of the expression architecture of breast cancers. Using gene expression driven in *cis* by CNAs, we have generated a new molecular taxonomy of breast cancer with 10 genomic driver-based subtypes termed Integrative Clusters. The samples used in this analysis were derived from the METABRIC cohort, which encompassed a large biobank of fresh frozen tumour samples collected across five major teaching hospitals in the UK and Canada ^4^.

Formalin-fixed paraffin-embedded (FFPE) tissue samples are more routinely collected and hence more representative of cancer in the general population. These FFPE archives are a valuable resource for molecular profiling in cancer research. Whilst the fixation process is essential to protect cellular morphology and protein expression, it is detrimental to nucleic acids and results in their chemical modification and degradation. As a result, extraction of DNA from FFPE tissues results in lower yields when compared to extraction from fresh frozen tissues. DNA extracted from FFPE works well for downstream applications using polymerase chain reaction (PCR), particularly for small size amplicons (less than 300 base pairs), but for other applications, including microarray based CGH, where efficient labelling of the DNA is dependent on its integrity, its use is more challenging. There have been several studies describing different methods for DNA extraction ^5^, quality control ^6,7^ labelling ^8^ and other optimisation protocols ^9^ to improve the performance of FFPE DNA on microarrays. In the past, we have tried to profile CNAs using FFPE DNA on microarrays with limited success. Only Illumina Infinium and Molecular Inversion Probe (MIP, Affymetrix) arrays yielded good results but these required good quality and at least 200ng of DNA ^10^.

Next generation sequencing has revolutionised cancer genomics. It is now relatively easy and inexpensive to sequence an entire genome. However, as with microarrays, the robustness of the results obtained are dependent on the quality of the input DNA. Two recent studies have demonstrated the feasibility of doing shallow whole-genome sequencing (sWGS) for CNA profiling using DNA extracted from FFPE tissue material ^11,12^ The first report used 250ng of DNA from FFPE tissues and a breast cancer cell line to produce libraries and developed an analytical method for sWGS. The second study compared several sequencing library production kits and reported generating successful sequencing libraries with low input DNA in a small number of FFPE samples.

Here we present extensive sWGS data generated from DNA extracted from FFPE breast cancer samples to describe steps to ensure successful libraries.

## Materials

### Specimen collection

FFPE tissue samples from invasive breast cancer patients diagnosed between 1997 and 2014 were obtained from several tumour repositories: Addenbrooke’s Hospital in Cambridge (n=62), a consortium of hospitals participating in clinical trials (GEICAM) in Spain (n=172), and Samsung Medical Center in South Korea (n=122). In some cases, we extracted DNA from adjacent normal (n=15) and DCIS (n=115) samples. Some of the clinical trials samples were biopsies taken at diagnosis (n=107) and/or surgery (n=106) where 41 are paired. All tumour samples were collected with informed patient consent and their use for genomics profiling had ethics approval from the institutional review board for each of the biobanks (Cambridge: REC ref 07/H0308/161; South Korea: 2014-10-041; Spain: NCT00432172 & NCT00841828). Detailed information on the sample cohort is collated in Table 1.

**Table 1:**
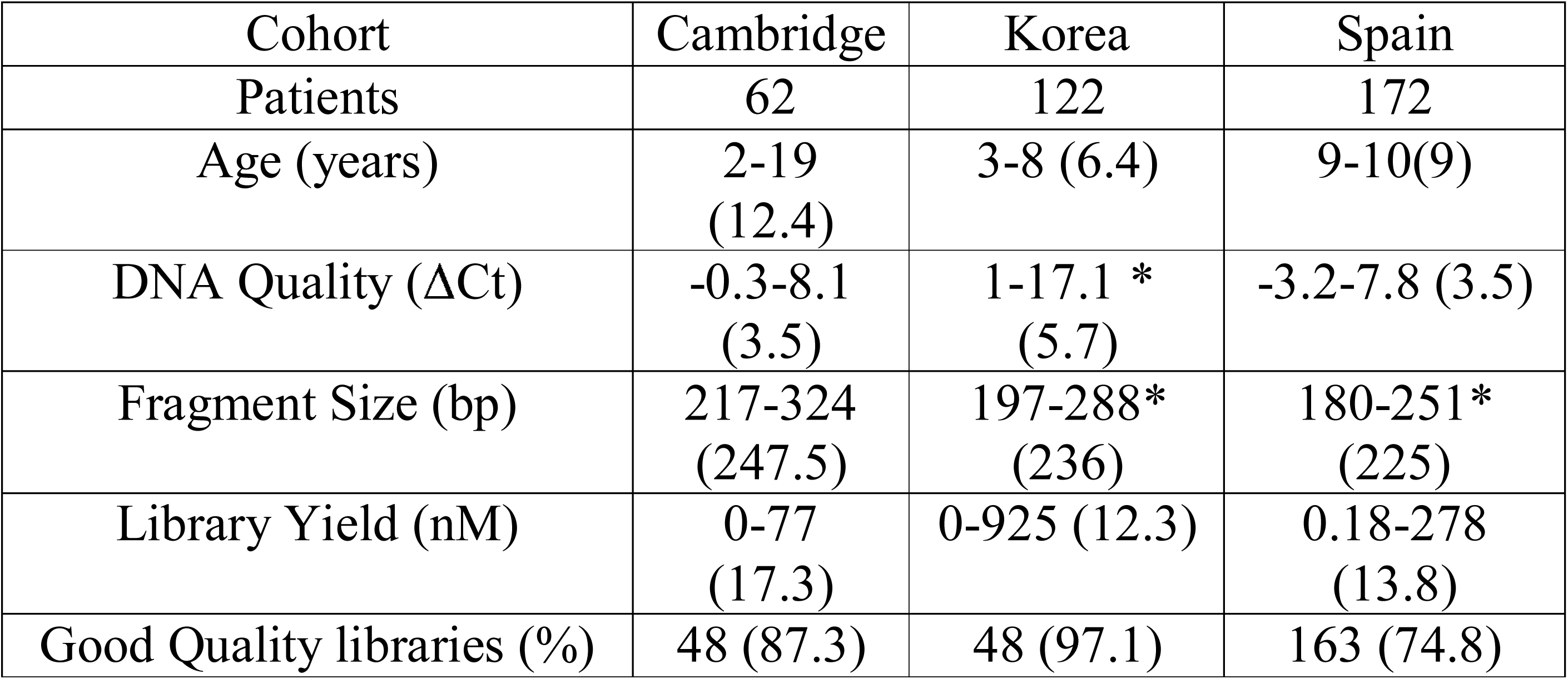
Description of the samples and libraries according to sites. Features of input DNA and libraries generated from FFPE blocks collected at three different sites. Data provided in minimum-maximum range and median in brackets. Age = years since blocks were generated. ΔCt = difference between the cycle threshold of test to the control template ACD1 provided in the kit. ng=nanogram, PCR= polymerase chain reaction, bp=base pairs, nM=nanomoles. * denotes the site where there is a significant difference to the index group (ie Cambridge).

### DNA extraction and quality control

DNA was extracted from either one mm cores punched from tissue blocks or from 10 x 30 micron sections (Cambridge and Korea), or 4-6x10 micron sections (Spain) from FFPE blocks, using Qiagen QIAmp DNeasy Kits (Qiagen, Germany) according to the manufacturer’s instructions. All DNA samples were quantified fluorometrically using the Qubit dsDNA High Sensitivity Quantification Reagent (ThermoFisher, USA). The DNA quality was assessed using Illumina’s FFPE QC kit, a quantitative PCR (q-PCR) assay. All test DNAs and the template control provided in the kit (ACD1) were diluted to 0.25ng/μl and PCR reactions set up in triplicate as per manufacturer’s instructions. DNA quality was quantified as the difference between the Ct (cycle threshold) value of the test FFPE-extracted DNA against the Ct value of the control DNA template.

### DNA Fragmentation

DNA samples of different concentrations (4-500ng) were diluted in water to a final volume of 15μl in Covaris microTUBE-15 8 strip tubes (Covaris, USA) and fragmented to an average size distribution of 150-180bp with Covaris LE220 Focused Ultrasonicator with Adaptive Focused Acoustics technology. The following parameters were used for shearing: Peak Incident Power: 180W; Duty Factor: 30%; Cycles per Burst: 50; with the fragmentation time: 250s for DNA with ΔCt <10, and 200s for DNA with ΔCt ≥10.

### Sequencing library generation

Sequencing libraries were generated using either the beta testing version of the Illumina FFPE TruSEQ kit (ILMN, libraries=45) or the Rubicon Genomics Thruplex DNASeq (RGT, libraries=446), as per manufacturer’s instructions. For four samples, we generated sequencing libraries using both kits to compare their performance (Supplementary Figure 1a-b). The sample metrics for both kits are presented in Supplementary Table 1.

**Figure 1.**
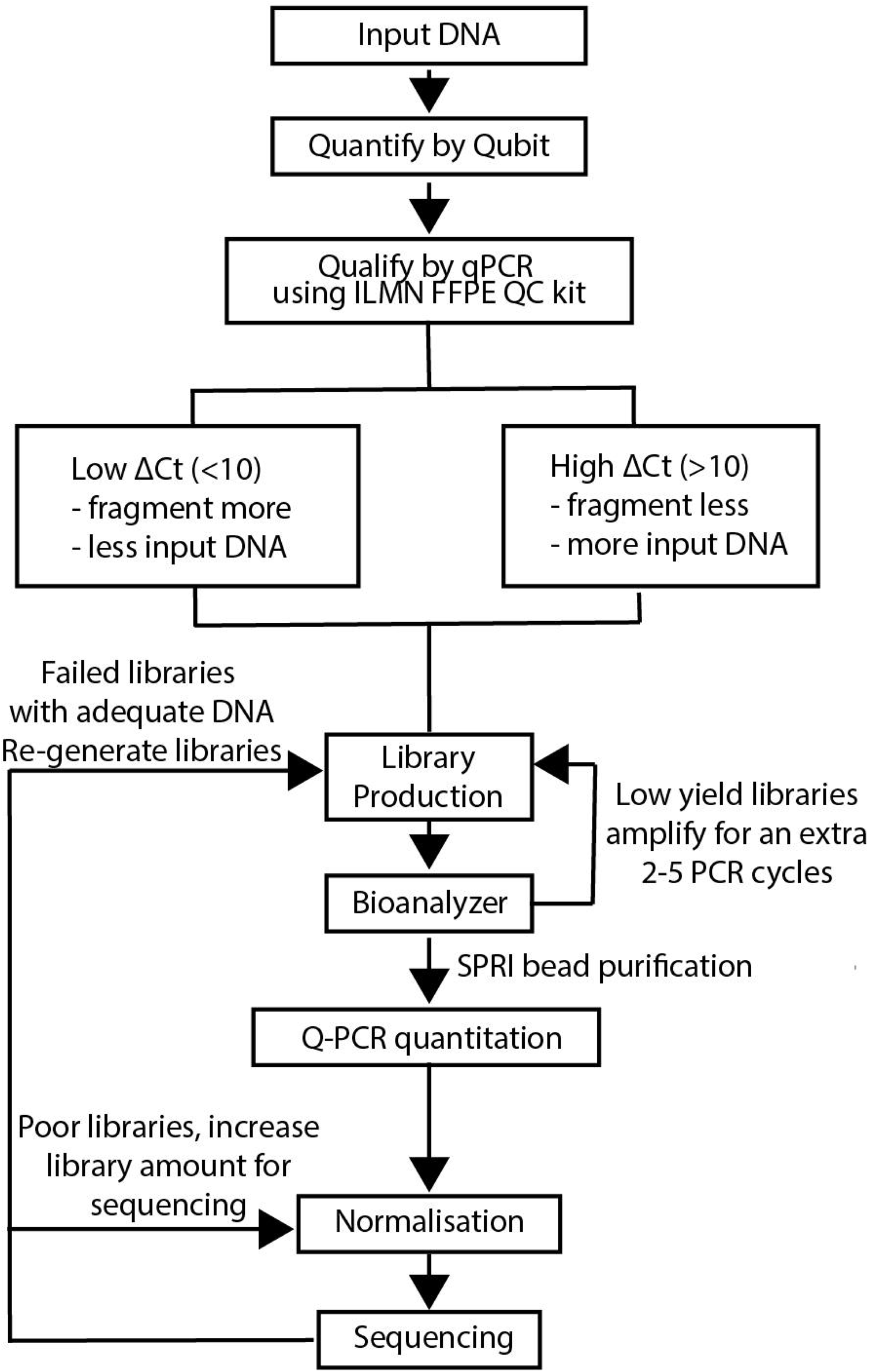
Overall Design. Schematic showing the workflow to ensure successful shallow whole genome sequencing (sWGS) libraries

The ILMN libraries were generated manually whilst RGT libraries were generated either on the Agilent Bravo (n=228) or manually (Spain, n=218). Final libraries were purified using magnetic beads (Agencourt SPRI beads, Becton Dickinson, USA) and eluted libraries were quantified using Kapa Library Quantification kit (Roche Life Technologies, USA). Fragment size distributions were analysed utilising a 2100 Bioanalyzer with a DNA High Sensitivity kit (Agilent Technologies, USA). Two nanomoles (nM) of each library were prepared and 48 samples were pooled in one lane for sequencing on a HiSeq4000 (Illumina, USA). The pools were re-quantified and normalised to 10 nM. Single end sequencing was conducted for 50 cycles, generating on average 4.3×10^8^ reads per lane.

### Bioinformatics

Alignment against the GRCh 37 assembly of the human genome was performed using BWA ver. 0.7.9^13^ or NovoAlign ver. 3.2.13 (NovoCraft, Malaysia). PCR and optical duplicates were identified using Picard tools (https://broadinstitute.github.io/picard) or Novosort (NovoCraft, Malaysia). Circular binary segmentation on the aligned files was performed in 100kb windows using the QDNAseq R package available on Bioconductor, which corrects for mappability and GC content ^11^. All statistical analyses were performed in R using the functions lm() for fitting linear models and t.test() for Welch two-sample t-test.

## Results

The majority of the FFPE samples available were core biopsies collected as part of a neoadjuvant clinical trial (GEICAM/2006-03, n=107) yielding low amounts of DNA (range=4 - 61ng, median 30ng). Therefore, to successfully generate libraries for CNA profiling using limited input DNA, we needed to understand how different variables could influence the quality of libraries and steps that can be taken to ensure good sequencing results (Figure 1).

### Assessment of the copy number plots

We examined the copy number plots by manual inspection and categorised them based on the variance in the CN data for each case into categories: “Very Good”, “Good”, “Intermediate” and “Poor” (Figure 2a). We also used QDNAseq ^11^ which calculates the expected (estimated from read depth) and measured (using read depth and influenced by DNA quality) standard deviation of the summarised reads, as a measure of variance. Both measures increased as the quality of library decreased and validated our categorisation of library quality (measured standard deviation shown in Figure 2b).

**Figure 2.**
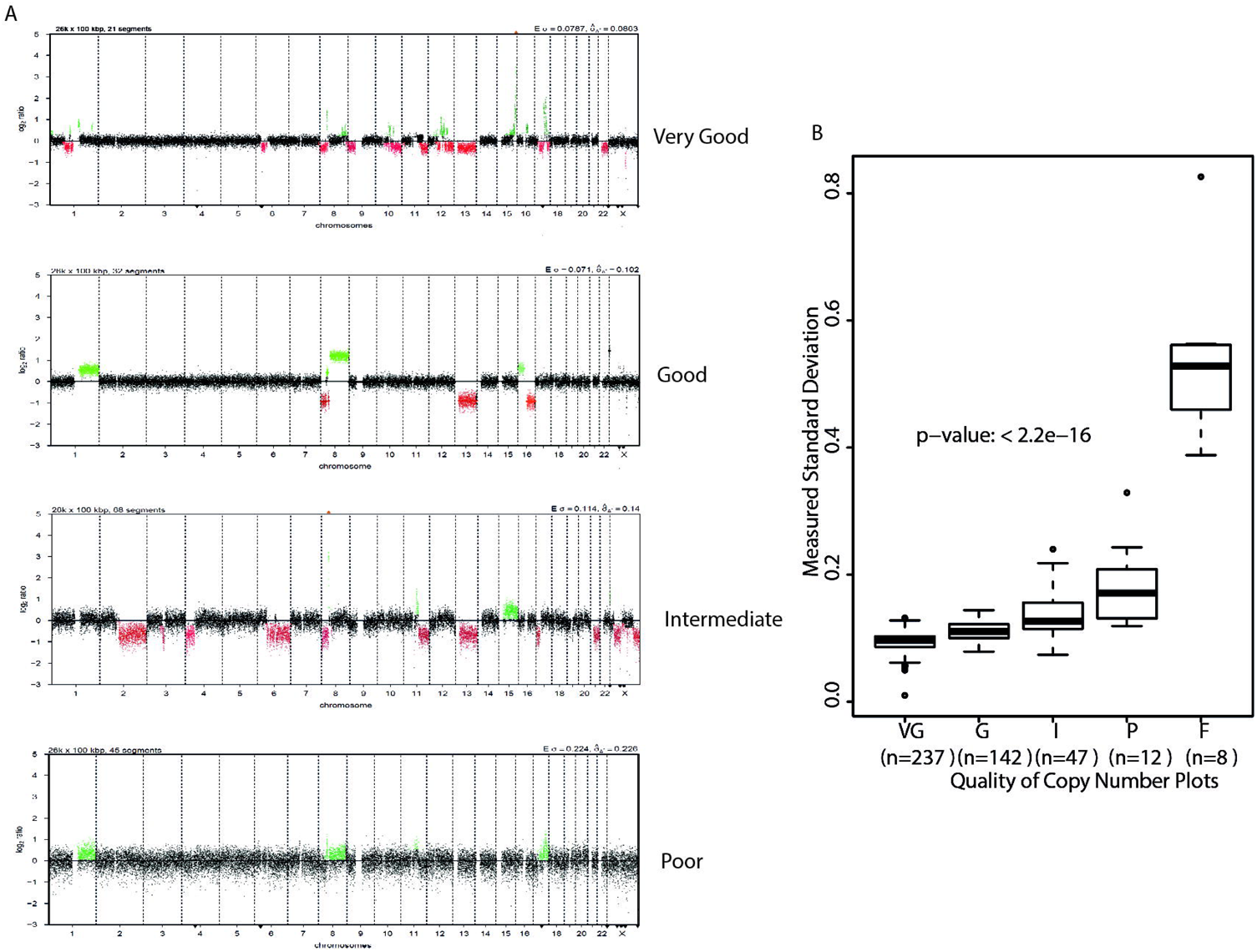
Categorisation of copy number profiles. A. Examples of QDNASEQ copy number plots scored as Very Good, Good, Intermediate and Poor. Failed libraries had very few reads and are not shown. Green dots represent regions of gains/amplifications and red dots represent regions of loss/deletion. B. Boxplots showing increasing measured standard deviations with decreasing libraries’qualities. Dots represent individual samples within each category. VG=very good, G=good, I=intermediate, P=poor, F=fail

### Assessment of different sequencing kits

We tested two kits (Illumina FFPE TruSEQ kit and Rubicon Genomics Thruplex DNASeq) using four FFPE samples to generate sequencing libraries and found comparable results (Supplementary Figure 1a-b). The CNA profiles obtained using DNA processed with the ILMN kit had less variance (noise) than those processed using the RGT kit however the ILMN libraries were generated using more input DNA (200-500ng (ILMN) versus 50ng (RGT)) and were sequenced deeper (average coverage 0.9X (ILMN) versus 0.08X (RGT)). For a more comparable evaluation, we down-sampled ILMN sequencing data to a similar read depth as RGT; this showed comparable copy number profile qualities between the two library preparation technologies.

In theory, increasing the sequencing depth should improve the copy number results by reducing the variance. We examined this by increasing the sequencing depth of 23 RGT kit libraries which had less reads (from 0.08X up to 0.15X) and found improvement in the data quality in 20 out of 23 libraries (examples shown in Supplementary Figure 2a). To examine the association between sequencing depth and variance, we down-sampled the number of reads (in steps of 1×10^6^ reads) for six libraries with high read counts (up to 24×10^6^ reads). We found a significant improvement in the quality of copy number plots with increasing number of reads (p<2.2e-16; Supplementary Figure 2b). It is interesting to note that the noise reduction levels off at approximately 7×10^6^ reads suggesting that increasing the read depth more than 7×10^6^ reads provides little benefit to variance reduction.

### Performance of sWGS for copy number profiling using the RGT kit

Due to the limited amount of DNA available for most samples, we chose the RGT kit as it required less input DNA due to fewer processing steps, in particular purifications. Sequencing libraries were generated from as little as 3.8ng of DNA, and out of 16 libraries prepared from less than 10ng of DNA, only one failed, 13 generated good quality CNA plots, and 2 generated intermediate quality CNA plots. Information for all the libraries generated are summarised in Supplementary Table 3.

### Recovery of under-performing RGT libraries

Eight (1.8%) libraries failed and 12 (2.7%) generated poor quality libraries out of 446 libraries. To recover some of these failed/poor samples, we prepared fresh libraries from samples with sufficient DNA (n=6) or repeated the sequencing using three-fold more library material for samples with insufficient DNA to generate new libraries (n=8). Thirteen of these new/re-sequenced libraries generated good quality data. The one repeat sample that failed was from the re-sequencing group. Consequently, only two out of 446 RGT libraries (taking into consideration the repeated libraries and re-sequencing) failed, resulting in a 99.5% success rate. Good sWGS data produced from 379/446 (84.9%) samples.

### Association between FFPE storage time, site, and sequencing quality

The FFPE samples were collected from three different tissue banks, spanning 20 years (Table 1). The effect of storage time on the DNA extracted was analysed (Figure 3). DNA from older FFPE blocks (>5years) was generally of poorer quality: higher ΔCt values, shorter fragment size, generating lower yield sequencing libraries. We compared the quality metrics for each banking site and found that overall FFPE samples from different sites were comparable (Table 1 and Supplementary Figure 3).

**Figure 3.**
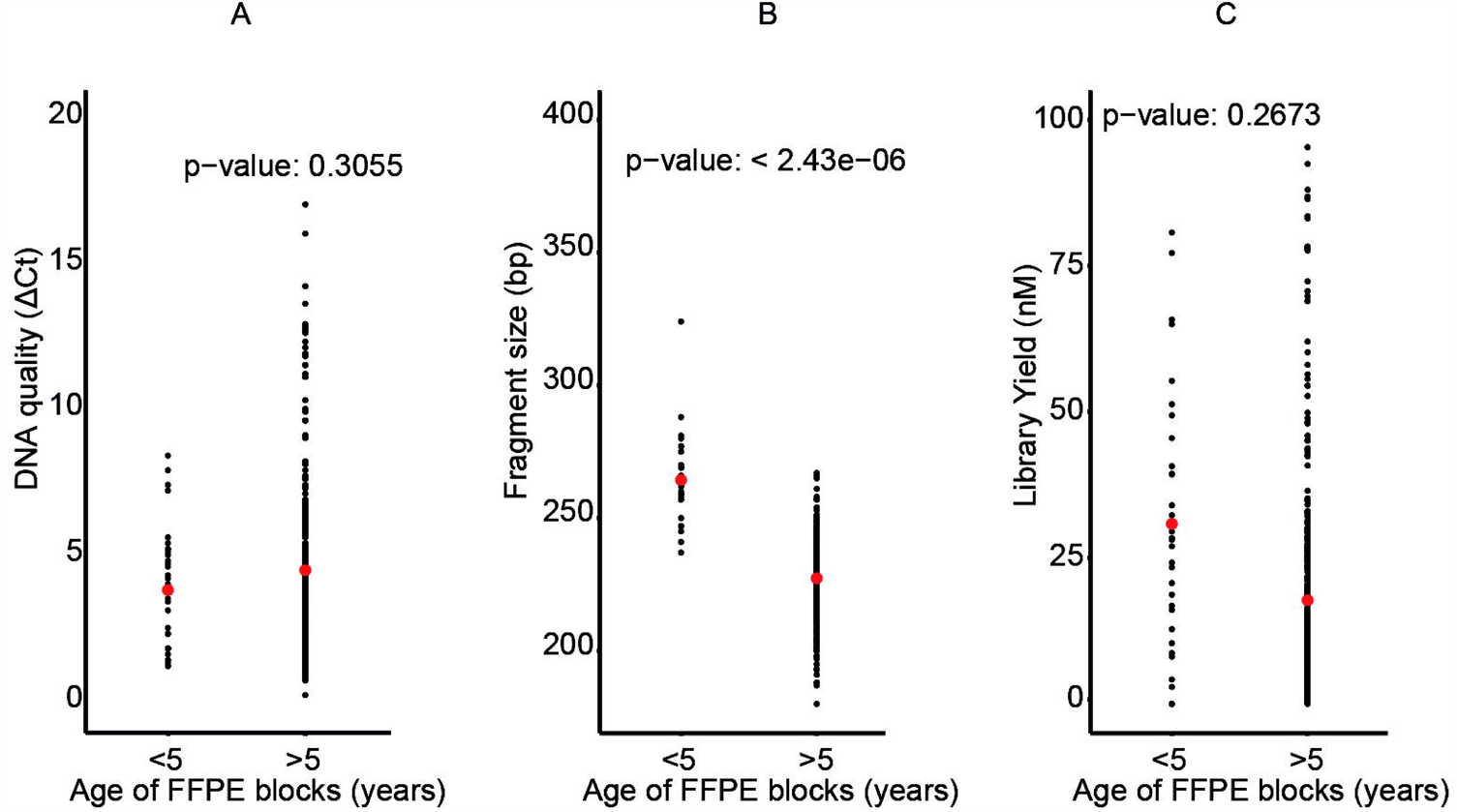
Features of input DNA and libraries generated from blocks less and more than five years. Dot plots represent the range (minimum-maximum) of observed values for each of the following categories and the red dot represents the median. A. The quality of input DNA inferred by ΔCt. B. Fragment sizes of the libraries in base pair C. The library yield in nanomoles

### Association between input DNA characteristics and sequencing library yield

We used the Illumina FFPE QC kit, a quantitative-PCR assay to estimate the quality of FFPE-extracted DNA. This assay measures the difference in Ct (cycle threshold) value of the test FFPE-extracted DNA against the Ct value of the control DNA template provided in the kit. Increasing ΔCt values indicate decreasing DNA quality with Illumina quality thresholds set at: ΔCt<1.5 denotes high quality (HQ), ΔCt<3.0 denotes medium quality (MQ), and ΔCt>3 denotes low quality (LQ) DNA. The Illumina DNA-input recommendations for sWGS are 50ng DNA with HQ DNA, 200ng with MQ DNA, and exclusion of LQ DNA. Using the ILMN kit, we could generate good quality sWGS using 50ng HQ and MQ DNA, and 200-500ng of LQ DNA. Unsurprisingly, for eight samples with paired libraries generated from 50ng and 200 or 500ng of input DNA using the ILMN kit, we found that the sequencing library yields generated with more DNA was significantly higher than when using only 50ng (p–value: 0.000265; Supplementary Figure 3a). This is an important consideration if these libraries were destined for downstream target enrichment assays for mutation detection that require 500ng of library material. Data from all the generated ILMN libraries (n=45) showed a library yield that averaged 5.6nM using 50ng FFPE-extracted DNA, which was significantly less than with libraries made with more input DNA (200ng: 23.6nM, Welch Two Sample t-test, p=5.24e^-06^; 500ng: 23.0nM, Welch Two Sample t-test, p=0.0121). There was no difference in library yield when using either 200 or 500ng of DNA (Welch Two Sample t-test, p=0.2401). This is probably due to the quality of the input DNA as libraries produced from 200ng of DNA had lower ΔCt values (better quality) than those using 500ng (Welch Two Sample t-test, p=0.0179, Supplementary Figure 4a-c).

Using the RGT kit, we found no correlation between amount of input DNA and sequencing library yield (r^2^= −0.002, p=0.81). This is probably due to the fewer library-washing steps using the RGT kit (six washing steps in the ILMN protocol versus one in RGT).

### Association between input DNA characteristics and sequencing library quality

Next we sought to determine if sequencing quality was influenced by the nature of the input DNA by looking at the proportion of samples from all quality (ΔCt) groups (Figure 4a), fragment sizes (Figure 4b) and different input groups (Figure 4c) in each of the sequencing quality categories. Reassuringly, we found no biases in sampling that contributed to the sequencing quality. In other words, each copy number plot quality group had samples from all DNA quality (ΔCt) groups, fragment sizes and input quantity groups, suggesting that we could generate good quality libraries from most of our FFPE DNA regardless of these features.

**Figure 4.**
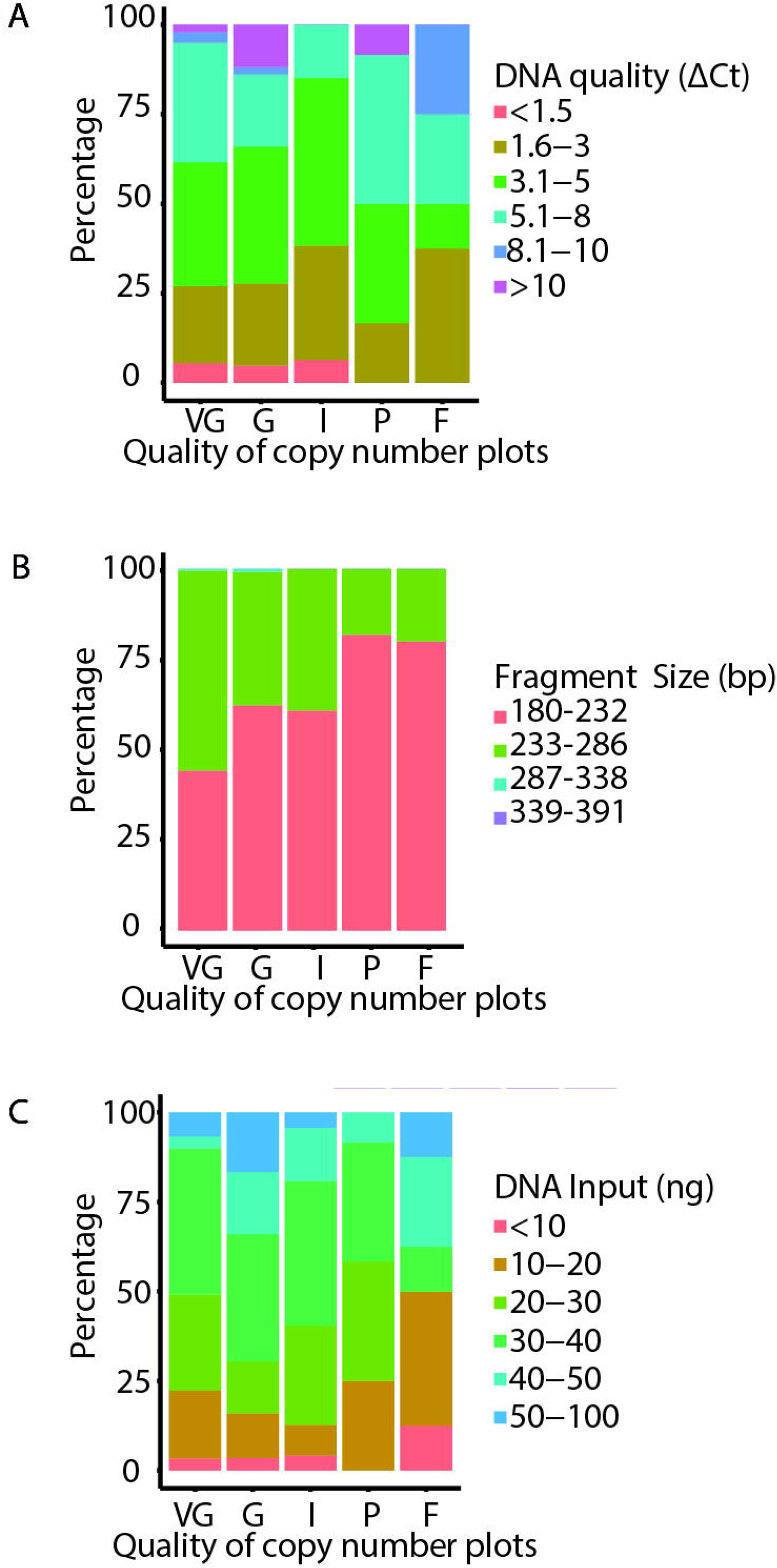
Measured standard deviations from the QDNASEQ copy number plots and associations with the quality of sequencing libraries. A. Bar charts showing proportion of samples with different input DNA quality (based on ΔCt) in each sequencing quality group. B. Bar charts showing proportion of samples from FFPE blocks of different fragment sizes in each sequencing quality group. C. Bar charts showing proportion of samples with different amount of input DNA in each sequencing quality group VG=very good, G=good, I=intermediate, P=poor, F=fail D.

Using our copy number output categorisation scoring, we examined if quality of the libraries (analysed as “all sequencing quality groups” versus “very good”) can be attributed to the different features of the input DNA and library yields (Table 2, Figure 5). We found that the quantity of template was only significantly different in the good quality libraries. Meanwhile, the quality of input DNA was significantly different in the intermediate libraries only when compared to the “very good” libraries. Therefore, the lesser quality sequencing libraries (I, P and F) cannot be attributed simply to either quantity or quality of the template DNA. The DNA fragment sizes, which should reflect the length of the template as the DNA was sheared under similar conditions, were found to be significantly different in all groups (progressively becoming shorter) except the failures. We found that low quality DNA was associated with shorter DNA fragments, lower library yield and higher number of unmapped reads but no association with the total number of unique reads aligned (Figure 6a-d). The recovery of most of the poor/failed libraries described previously, was achieved by either repeating the library generation or re-sequencing to generate more reads. Consequently, we suspect the poor/failed libraries could be due to a loss of DNA during the purification steps or that the Q-PCR quantification of the libraries prior to normalisation, over-estimated the library concentration resulting in inadequate amount of library being used for sequencing. This would explain why by simply increasing the quantity of libraries for sequencing and reducing the number of samples in a single pool, ensured adequate read counts and successful sequencing.

**Table 2:** Description of the libraries grouped by quality. Features of input DNA and libraries for the different categories of copy number data. Data provided in minimum-maximum range and median values in brackets. ΔCt = difference between the cycle threshold of test to the control template ACD1 provided in the kit, ng=nanogram, bp=base pairs, nM=nanomoles, SD=standard deviation. * denotes the site where there is a significant difference to the index group (ie Cambridge).

| Quality | DNA Input (ng) | DNA Quality (ΔCt) | Library Yield (nM) | Fragment Size (bp) | Observed SD |
| --- | --- | --- | --- | --- | --- |
| VG | 3.8-100<br>(30.3) | -0.3-12.7<br>(4.4) | 0.28-245.1<br>(16.1) | 180-288<br>(235) | 0.01-0.132<br>(0.097) |
| G | 4.3-107<br>(35.9)* | 1-17.1<br>(4.2) | 0.18-139.7<br>(11.3) | 195-324<br>(229)* | 0.0789-0.144<br>(0.1105)* |
| I | 4-59<br>(33.1) | -3.2-6.3<br>(3.5)* | 1.06-924.8<br>(16.0)* | 191-258<br>(231)* | 0.0739-0.24<br>(0.127)* |
| P | 14.7-50<br>(28.7) | 2-11.3<br>(4.9) | 0-55.63<br>(8.7) | 187-243<br>(219)* | 0.119-0.329<br>(0.171)* |
| F | 4.8-51.1<br>(25.2) | 1.9-8.2<br>(4.9) | 0-179.6<br>(8.2) | 204-244<br>(225) | 0.388-0.826<br>(0.528)* |

**Figure 5.**
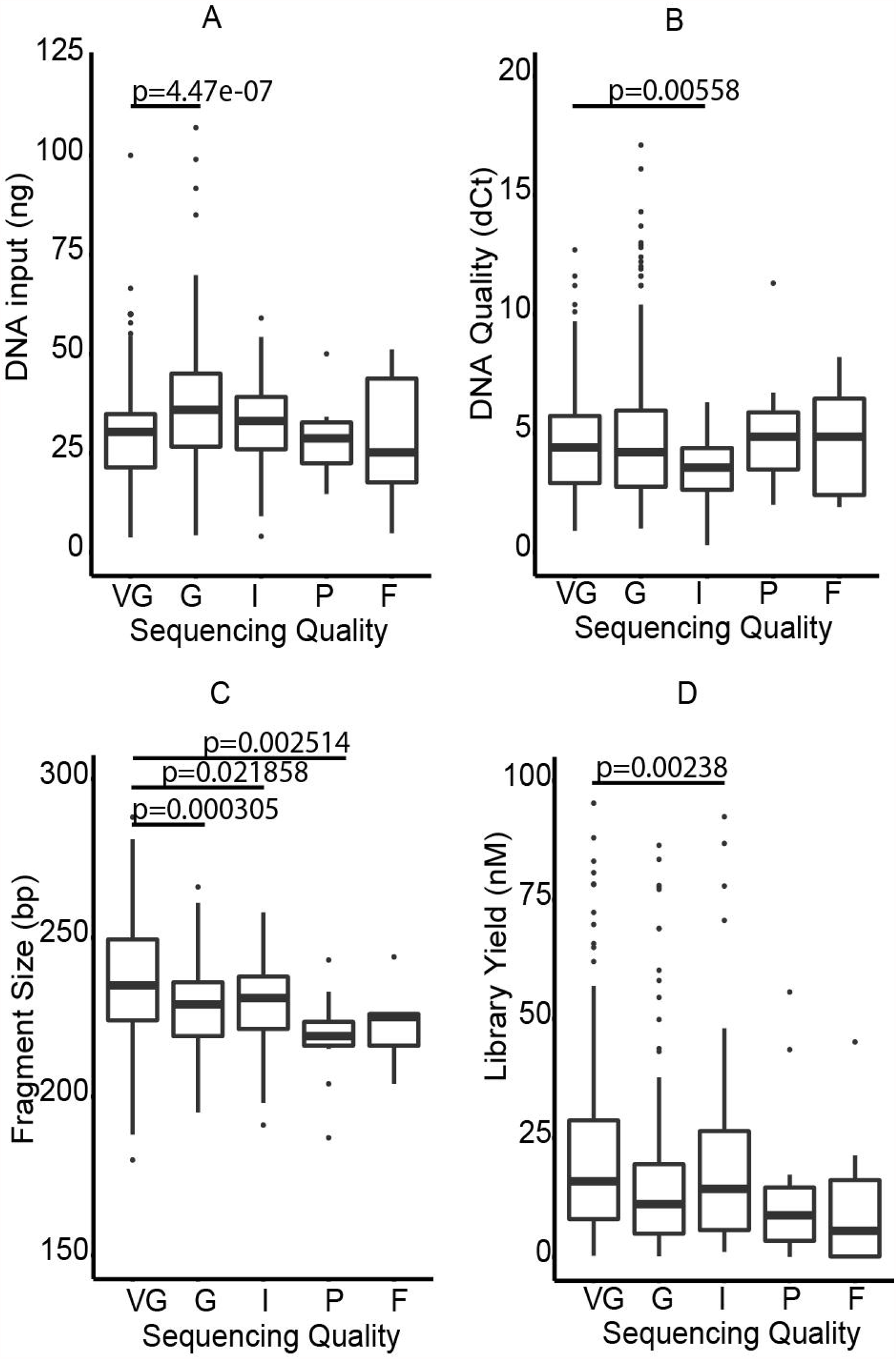
Features of the sequencing libraries. Boxplots showing different features of input DNA and library yield relative to the different library qualities. A. Input DNA B. Quality of input DNA inferred from ΔCt C. Fragment size of libraries D. Library yield VG=very good, G=good, I=intermediate, P=poor, F=fail

**Figure 6.**
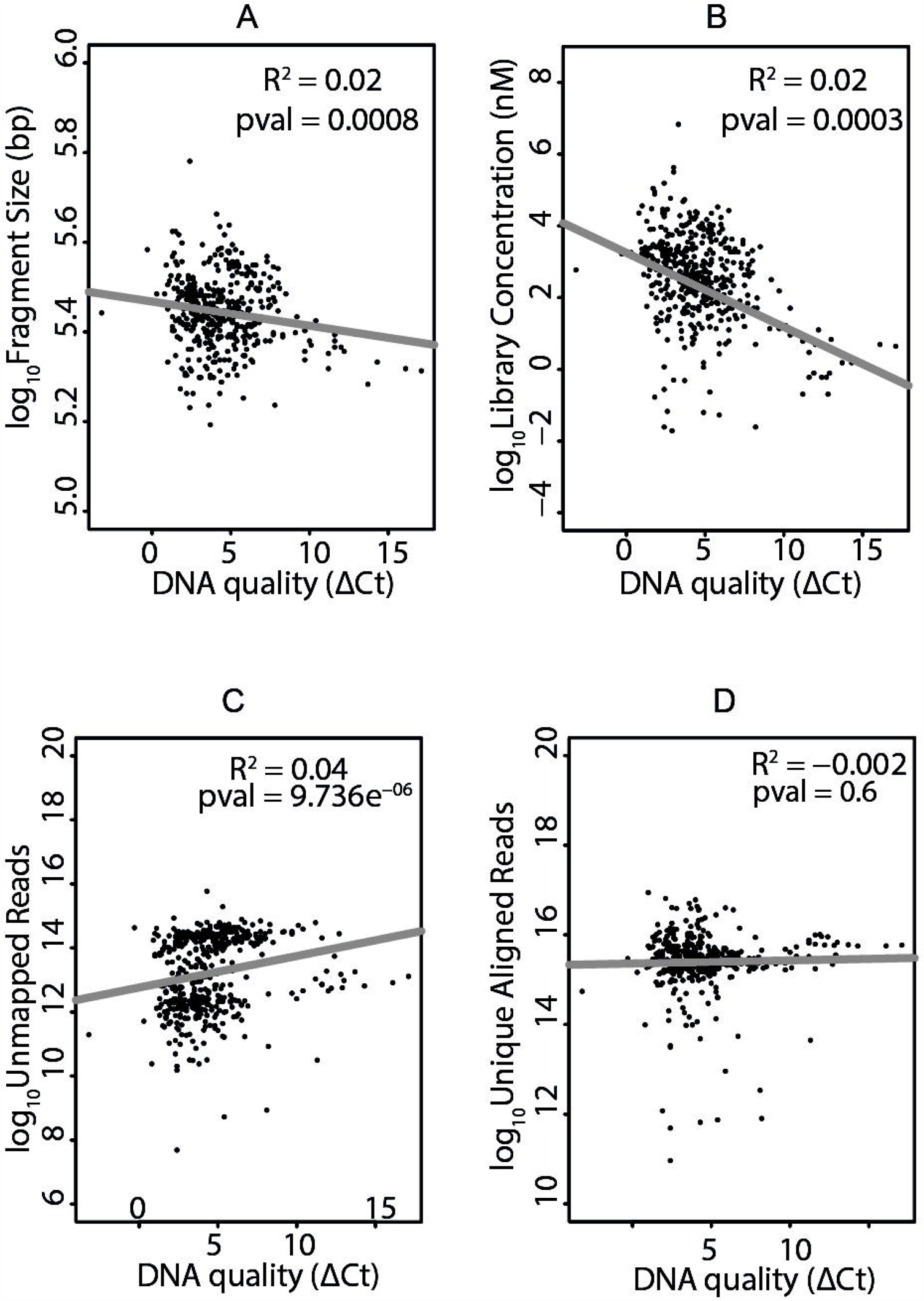
Effect of input DNA quality. Scatterplots showing the association between quality of input DNA with different features of the sequencing libraries. A. Fragment size of libraries B. Library yield C. Unmapped Reads D. Unique aligned reads

## Discussion

In this study, we have looked at the effect that quantity and quality of DNA from FFPE tissues has on successful sWGS library preparation for CN profiling of human breast cancers. Both the quantity and quality of DNA have always been an important consideration for sample selection and in deciding which genomic application to use. For example, microarrays require 100ng – 2.5μg of DNA depending on the resolution of the arrays whereas PCR based methods require only 10ng of DNA. In our hands, we have not had much success in obtaining CN data with DNA extracted from FFPE DNA using microarrays, especially when the extracted DNAs are more fragmented and of lower quality (judging from absorbance ratios of 260nm to 280 nm and multiplex PCR for quality control).

Here we have robustly shown that we can generate CN data from virtually all archived FFPE samples using sWGS. We show good CN profiling data irrespective of the quality of input DNA, as inferred by whether it can be amplified with Q-PCR (ΔCt). Previous work has extensively tested the utility of FFPE DNA for mutation analysis ^14-18^ but to date no comprehensive study has shown its use for CN profiling. Since many human cancer types, including breast and ovarian cancers, are driven mostly by CNA (C-class) rather than point mutations or indels (M-class), we believe more effort should be focussed on characterizing the copy number landscapes of these cancers ^3^. We found sWGS to be very robust in generating these CN profiles, independently of the kits used, quantity and quality of DNA. sWGS is also significantly cheaper (~50%) than microarray-based methods (Supplementary Table 2).

Another advantage of generating sWGS libraries is the ability to use the same library for targeted sequence enrichment to identify mutations. There have been other methods reported for CN profiling using DNA extracted from FFPE samples but these methods do not generate sequencing libraries that can then be used for target enrichment and sequencing ^19^ or if they do, are expensive ^20^. In addition, sWGS will also serve as a quality control for the libraries, given its relative low cost when compared to that of generating targeted sequencing libraries. Only libraries that generate good CN profiles should be used for target enrichment and mutation detection^21^. Whilst we haven’t performed target enrichment on our FFPE libraries, we expect the performance of these FFPE libraries for mutation analysis to be similar to that of published data, including known artefacts caused by formalin-based fixation effects on the DNA template ^15,17,22-24^

In summary, we have shown that sWGS is a robust and cost-effective method for obtaining good quality CN data from FFPE cancer samples, irrespective of the DNA quality and quantity used. In the case of breast cancer, CN profiles can be used to stratify breast cancers into one of the 10 Integrative Clusters ^25^, reiterating the importance of FFPE tumour archives. The methods described here are also of relevance to other cancers, e.g. ovarian cancers where CN profiling is essential to characterise their genomic landscapes.

## Acknowledgements

This sequencing project was funded by CRUK. We thank the Genomics, Histopathology, and Biorepository Core Facilities at the Cancer Research UK Cambridge Institute; the Addenbrooke’s Human Research Tissue Bank (supported by the National Institute for Health Research Cambridge Biomedical Research Centre). We thank all the patients who donated tissue and the associated pseudo-anonymized clinical data for this project.

